# Prevalent Integration of both Xenotropic and Ecotropic Murine Leukemia Virus in the Human Genome of Xenografts

**DOI:** 10.64898/2026.11.09.764062

**Authors:** Xiaoyu Zhuo, Noah L. Basri, Wesley N. Saintilnord, Xiaoyun Xing, Ting Wang

## Abstract

Humanized mouse models are widely used in medical research to represent human conditions (1, 2), but their fidelity could be compromised by mouse-originated infections. However, their susceptibility to mouse pathogens are rarely questioned (3). Here we screened 1,619 publicly available genomic sequencing runs from patient-derived xenograft (PDX) models for the integration of murine leukemia virus (MLV), a rodent retrovirus exists in the mouse genome as an endogenous retrovirus (4), within the human genome. We discovered substantial enrichment of human-MLV chimeric DNA in approximately 25% of them. Analyzing the human-MLV junctions, we found about 90% of them bearing the unmistakable hallmarks of retroviral integration, including the canonical MLV boundaries and the distinctive 4-bp target site duplications (5, 6). It was also assumed that mouse strains without the highly infectious *Bxv1* provirus are safe from MLV contamination in xenografts (7, 8). However, we found many PDX models derived from *Bxv1*- strains, including the widely used NSG strain, are also contaminated by the supposedly rodent-specific ecotropic MLV (E-MLV). We further confirmed a complete E-MLV integrated into the human genome by both HiFi and sanger sequencing. Our discovery raises concerns that mouse-to-human retroviral transmission and integration is more common and less restrictive than previously appreciated. They support examination of MLV integration as a feature of humanized mouse models with implications on the fidelity of these models in recapitulating human cells and tissues.

Our discovery also has underscored the importance of comprehensively inactivation of porcine endogenous retroviruses for the clinical application of xenotransplatation (9, 10).

**Significance Statement:** Humanized mouse models are widely used to study human diseases, but researchers often assume these models are rarely infected and therefore are not affected by rodent viruses. We analyzed 1,619 publicly available sequencing datasets from patient-derived xenograft models to search for integrated murine leukemia virus (MLV) in the human genome. We found strong evidence of retroviral integration in about 25% of them, including the expected retroviral integration signatures and a complete copy of ecotropic MLV integrated into human DNA. This demonstrated that mouse-derived retroviruses can enter human cells and integrate into human genome more often than previously believed. The results imply that such viral integration could alter the biology of humanized mouse models and should be carefully evaluated.

## Introduction

Retroviruses integrate their genome into the host’s chromosomes during replication (5). Infection and subsequent integration into host germline cells can enable vertical transmission to progeny as well as continued horizontal infectious propagation. Murine leukemia virus (MLV) is one such mouse endogenous retrovirus (mERV) that is present in the genomes of all commonly used mouse strains and retains the capacity to produce infectious viral particles. MLV is broadly classified into three types: Ecotropic MLV (E- MLV), which can only infect mouse and rat cells; Xenotropic MLV (X-MLV), which can infect other mammal cells but not mouse cells; and Polytropic MLV (P-MLV), which can infect cells from a wide range of mammals including mice (11).

Considerable efforts to uncover potential cross-species MLV infection in various human patients have been previously reported. From 2006 to 2015, hundreds of publications described the association between Xenotropic Murine leukemia virus–Related Virus (XMRV) and prostate cancer, myalgic encephalomyelitis/chronic fatigue syndrome (ME/CFS) and immunosuppression following bone marrow or solid organ transplantation (12). However, upon careful examination, most of these studies can be attributed to contamination or data fabrication. Consequently, despite being detected in 22Rv1 after years of serial passages, XMRV infection in humans and its association with disease states have been thoroughly refuted (12–16). Following the identification of XMRV, several studies reported X-MLV infection in various human cells (8, 17–22).

However, due to the experimental confounding and irreproducibility that plagued pathology studies of XMRV integration, the presence of murine DNA in xenografted samples continues to be regarded with skepticism by many scientists. Indeed, Bock et al. concluded that “mERV integration into the genome of human cells is unlikely or at least a very rare event,” after carefully measuring MLV levels from various tissues (7).

The presence of multi-system physiology and technical reproducibility have established murine models as an invaluable resource in studies of metabolism, immunology, neuroscience and oncology. Particularly in cancer research, patient-derived xenograft (PDX) models have been widely used to model intrinsic tumor features, such as cellular heterogeneity, drug sensitivity, mechanisms of evolved resistance, and other properties with higher fidelity than commercial cell lines (23). In 2016, the National Cancer Institute (NCI) made a significant shift by retiring the well-known NCI-60 panel of cancer cell lines in favor of hundreds of PDX models (24). Today, numerous publicly and privately funded PDX registries have been established, representing thousands of diverse tumors (1). They serve as valuable resources for cancer researchers worldwide, facilitating collaboration and advancing cancer research.

With the rapid advancement of disease mouse models including PDX models, the possibility of MLV infection has often been overlooked. Cross-species retroviral infection can introduce new genetic material into engrafted human cells, raising questions about its consequences for xenograft biology. Failure to accurately characterize the frequency and degree of them in xenograft studies invites potential experimental confounders and opportunities for misinterpretation of biological phenomena under study. For instance, mERVs natively contribute alternative transcription initiation and transcription factor binding sites, enhancers, topologically associating domain boundaries and protein- coding sequences, all of which can alter physiology in a human host cell in as yet unappreciated ways (25–28).

Here, we present Kmeric: a rapid k-mer-based tool for identifying human-virus chimeric reads in paired-end short-read sequencing data. Using Kmeric to screen publicly available PDX genomic reads for MLV integration, we discover approximately 1 in 4 sequencing files generated from PDX models are infected by MLV, and we report the first documented evidence of cross-species E-MLV integration in the genome of human cells. Our findings challenge the notion that E-MLV cannot infect human, and we propose consideration of unexpectedly widespread MLV integration as a potential confounder in mouse xenograft studies.

## Results

### Human-MLV chimeric reads are highly enriched in PDX DNA samples

Since murine cells are present in PDX models, the mere presence of MLV sequences is insufficient to infer MLV infection in human cells. We thus set to specifically identify MLV integration in the human genome using MLV-human chimeric reads from PDX-derived genomic DNA libraries catalogued in the National Center for Biotechnology Information (NCBI) Sequence Read Archive (SRA) (29, 30). In total, we identified 1,619 paired-end, genomic DNA sequencing runs (Whole Genome Sequencing, ChIP-seq, ATAC-seq, etc.) generated from PDX models containing 121 trillion pairs of reads (Table S1).

Applying the widely used k-mer classifier Kraken2, reads were categorized as human- only, mouse-only or human-mouse chimeric, wherein one end is classified as human and the other mouse (31). We further excluded low-complexity and low-quality reads (32, 33) and aligned the remaining chimeric reads to the human and mouse reference genome simultaneously. Finally, we selected chimeric reads where each end can be confidently aligned to either human or mouse reference genome as true human-mouse chimeric reads (34) (Materials and Methods) (Fig. 1A).

**Fig. 1:**
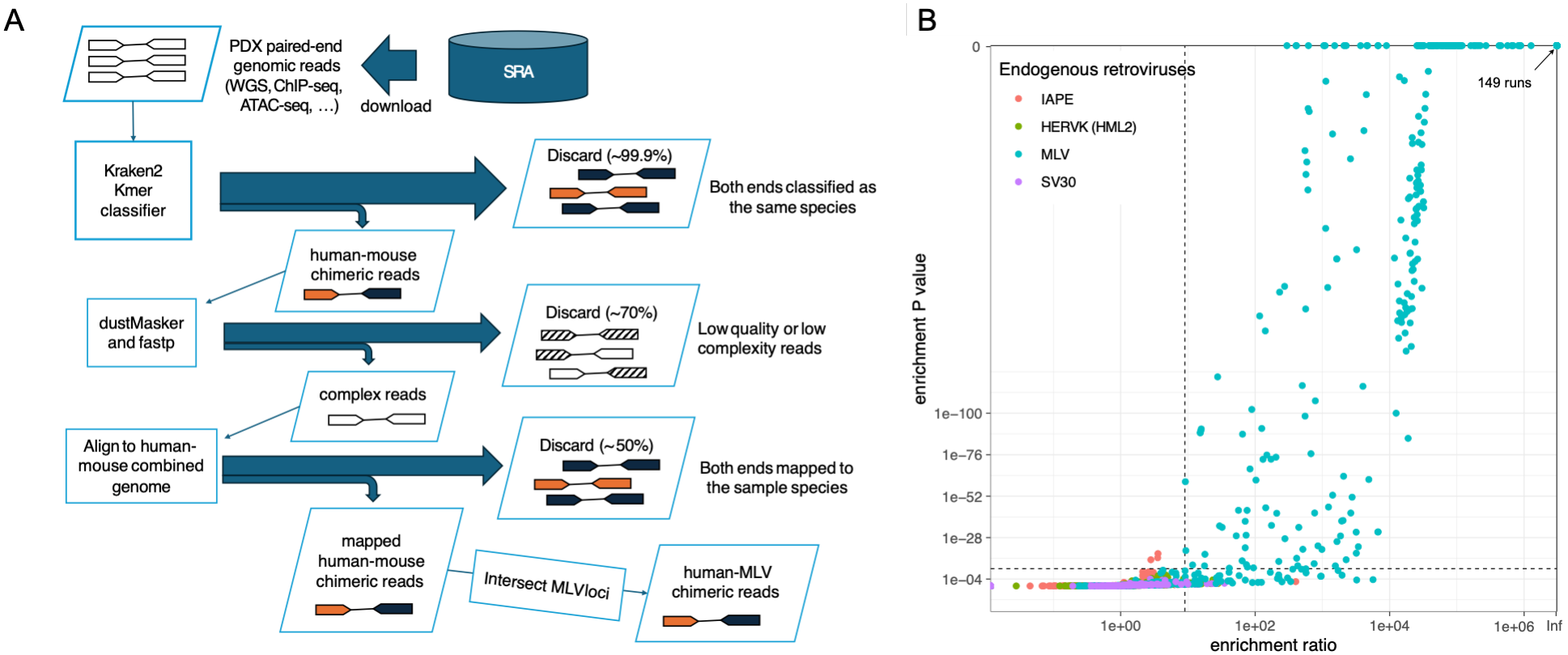
The enrichment of MLV-human chimeric reads. (**A**) Schematic of kmeric pipeline to extract human-mouse and human-MLV chimeric reads from paired-end PDX genomic reads. We downloaded sequencing raw reads from SRA, and classified them into human reads, mouse reads, or human-mouse chimeric reads using Kraken2. After filtering out human reads and mouse reads, we further filtered out low complexity and low quality reads by applying dustMasker and fastp. At last, we aligned the remaining reads to a combined human-mouse genome using bwa, and only kept paired-end reads with one end aligned to the human genome and the other end aligned to the mouse genome. (**B**) Fisher’s exact test enrichment ratio (X-axis) and one-tailed P-value (Y- axis) of MLV (RLTR4_mm), IAP (IAPEY), SV30 (RLTR6_Mm) and HERVK(HML2) (LTR5_Hs) in all human-mouse chimeric reads. Fisher’s exact tests were conducted using BEDTools per sequencing run. The enrichment cut-off lines (enrichment ratio > 9 and P value < 1e-09) were shown as dotted lines.

Observation of sequencing artifacts containing chimeric sequences during high- throughput sequencing is not uncommon (35). Collectively, all copies of the long terminal repeats (LTRs) of MLV (RLTR4_Mm) represent approximately 100KB within the roughly 3GB mouse reference genome; therefore, we posited a null distribution of ∼0.003% of human-mouse chimeric reads containing MLV if chimeric reads were randomly generated from the genome. However, we observed that human-MLV chimeras were substantially overrepresented among human-mouse chimeric reads in 397 out of the 1,619 (∼25%) PDX sequencing runs (Fig. 1B). Notably, the enrichment in 236 sequencing runs achieved a P-value of 0 (the upper boundary in the panel), and among them, 149 also reached an "infinite" enrichment ratio (upper-right corner) applying Fisher’s exact test implemented in BEDTools (36).

To assess whether this enrichment is generalizable across other active/polymorphic ERVs, we performed the same enrichment calculation for mouse IAPEY and human HERVK-HML2. We also calculated the enrichment of VL30, a non-autonomous mERV that can be packaged into MLV particles (37). However, we did not observe similar enrichment of human-IAPEY, human-VL30, or mouse-HERVK chimeric reads in the sequencing runs (Fig. 1B). These data indicate the contribution of MLV to human- mouse chimeric reads is significantly higher than what would be expected based on its contribution to the mouse genome and strongly suggests that these human-MLV chimeric reads represent true integrations rather than sequencing artifacts.

### The majority of human-MLV chimeric reads carry hallmarks of MLV integration

We extracted 252,187 human-MLV chimeric read pairs from the 397 sequencing runs where the human-MLV chimeric reads were enriched, and we grouped them into 27 samples based on the metadata from SRA (Table S2). Since a canonical MLV provirus in the genome starts and ends with a palindromic sequence: 5’-TGAAA …MLV… TTTCA-3’ (5), we expect to find human genomic sequences directly preceding the 5’- TGAAA-3’ motif in reads derived from human-MLV junctions (Fig. 2A). Accordingly, 93% of the 100,358 chimeric reads from MLV-enriched samples that have a soft-clipped overhang while aligning to the human reference genome matched such pattern (Fig. 2B). Also consistently, all human-mouse split-alignment reads piled up to both edges of MLV when we aligned them to the LTR of MLV (Fig. 2C). This observation suggests the majority of human-MLV chimeric reads from the 397 MLV-enriched runs perfectly match the boundary of a typical integration site of MLV, further signifying MLV integration sites in human xenografted tissues.

**Fig. 2:**
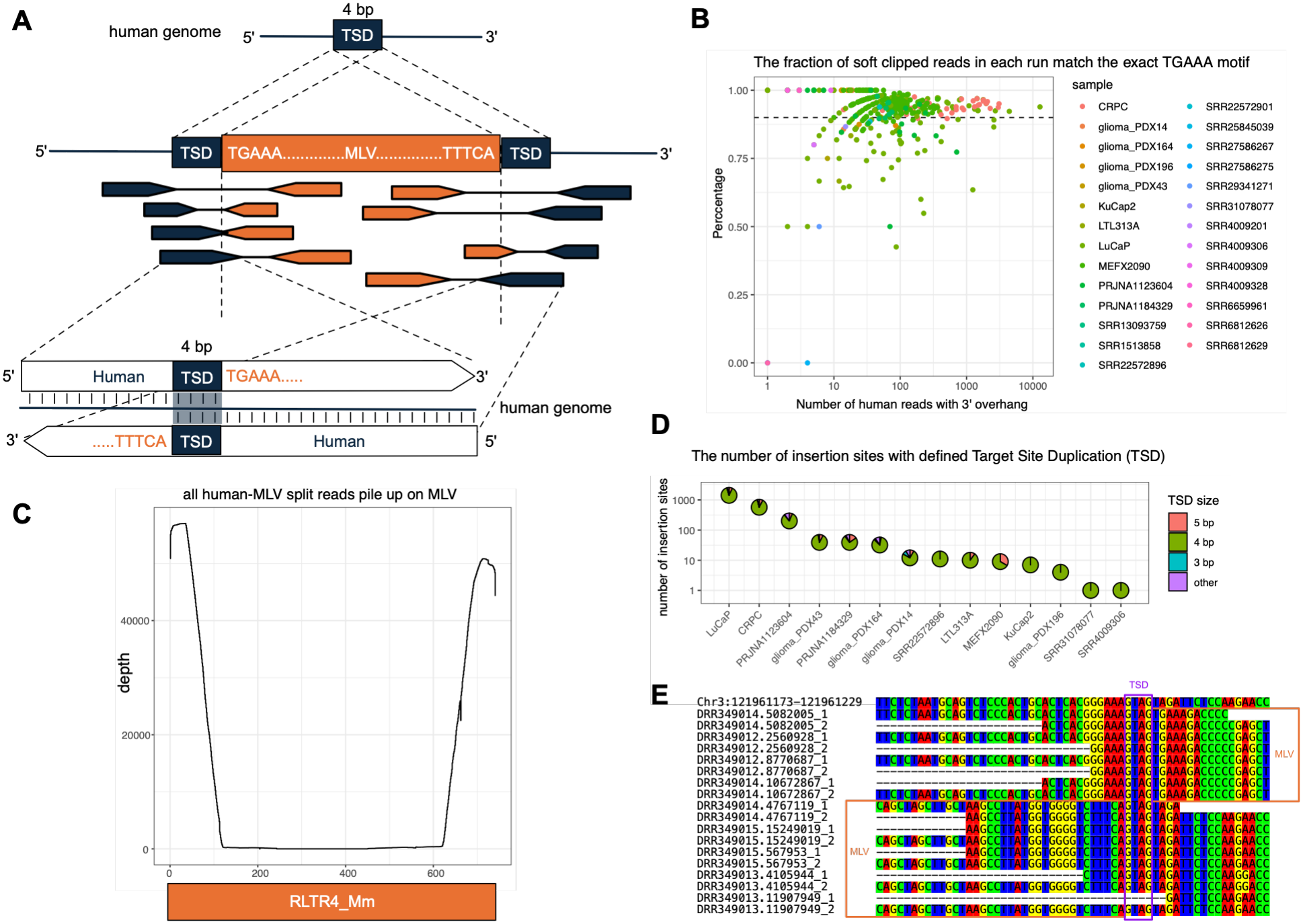
The characteristics of MLV integration. (**A**) Schematic of sequencing features of MLV integrations. Following retroviral integration, Target Site Duplications (TSDs) were created on the host genome (typically 4bp in the case of MLV). The MLV provirus in the genome begins and ends with the palindromic sequence 5’-TGAAA-3’. The bottom two illustrated human-MLV chimeric reads spanning the human-MLV boundaries were shown as alignment to the human reference genome with a 3’ end overhang. The TSD can be found as the overlap between them in the shaded area. (**B**) The fraction of human-MLV chimeric reads that begins the MLV part with 5’-TGAAA-3’ on the 5’ end. All 1,619 sequencing runs with human-MLV chimeric enriched were plotted, colored by their sample. X-axis represents the number of human-MLV chimeric reads in the sequencing run, Y-axis represents the fraction of all human-MLV chimeric that begins the MLV sequence with 5’-TGAAA-3’. (**C**) The mouse part of human-MLV split alignment reads pile up at the boundaries of the full-length MLV RLTR4_Mm. (**D**) The size of all identified TSD. The number and size of all TSDs in the 14 samples with TSDs identified. X-axis lists the 14 samples, Y-axis shows the number of insertion sites in each sample. The pie-chart in the grid shows the sizes of TSDs of insertion sites. (**E**) An example of MLV integration site at Chr3:121961208-121961212 in the human reference genome CHM13 with the TSD sequence GTAG from prostate cancer PDX KuCap2 in study DRP008380. The first row shows the human reference genome CHM13 from Chr3:121961173-121961229 and the following rows showed parts of reads that supported the insertion. Four pairs of ChIP-seq reads (DRR349014.5082005, DRR349012.2560928, DRR349012.8770687, and DRR349014.10672867) support the upstream insertion boundary and 5 pairs (DRR349014.4767119, DRR349015.15249019, DRR349015.567953, DRR349013.4105944, and DRR349013.11907949) support the downstream boundary. They were constructed with a small insertion size, and both ends of the 9 pairs overlap. All but one end (DRR349013.11907949_1) covered the human-MLV boundary.

Another characteristic of retroviral integration is the target site duplication (TSD), and the typical TSD of MLV is 4 base pairs (5, 6). If insertion sites can be identified with supporting reads at both the upstream and downstream boundaries, we can determine the presence and the size of TSD from the overlap between upstream and downstream reads aligned to the human reference genome (Fig. 2A). In 14 of the 27 MLV-enriched samples, we discovered 2,203 distinct MLV insertion sites that were supported by TSDs (Table S3). Among these, 88% were 4 bp long, while 7% were 5 bp long (Fig. 2D). The majority of the TSDs we identified matched the expected length of the canonical MLV insertion (5). Notably, only LuCaP from PRJNA657288, PRJNA679976 and PRJNA998718 and Castration-Resistant Prostate Cancer (CPRC) from PRJNA777879 shared 135 MLV insertion sites (Table S3), suggesting that these two samples originated from the same MLV-infected tumor, while all the others were independently infected by MLV.

We displayed one MLV insertion site with its supporting reads in Fig. 2E. Specifically, at Chr3:121961208-121961212 in the human reference genome CHM13, an MLV insertion was identified in prostate cancer PDX KuCap2. This event was supported by nine pairs of ChIP-seq reads from the study DRP008380 (38). Notably, both ends of each pair of reads overlap due to short genomic fragments in the DNA library. The presence of well- supported TSDs, a definitive hallmark of retroviral integration, provided concrete evidence that our observations indeed represent the integrations of MLV within the human genome.

### Possible integration of both X-MLV and E-MLV into the human genome

Human-MLV chimeric reads can be mapped to any of the approximately 100 MLV provirus or solo LTR loci in the mouse reference genome (C57BL/6J strain). Despite the high polymorphism of the MLV insertions among strains and the use of different mouse strains for xenograft, identifying which MLV on the mouse reference genome aligns best with each chimeric read can provide valuable insights into which MLV types are contributing to the infection and integration in the human genomes.

For each of the 27 samples, we piled up all the chimeric reads on the mouse reference genome (Fig. 3). Generally, chimeric reads obtained from different experiments exhibit distinct pileup peak profiles. However, we found samples share similar profiles if they were xenografted from the same mouse strain. Specifically, most of the human-MLV chimeric reads that were xenografted on SCID and BALB/c strains piled up on the *Bxv1* provirus locus on Chromosome 1. Our observation suggested that these samples could be infected by the highly active X-MLV *Bxv1*. This is consistent with the presence of *Bxv1* in these strains (8, 11) and the observation that most of the X-MLV contamination in human cells likely originated from *Bxv1* (17, 20–22).

**Fig. 3:**
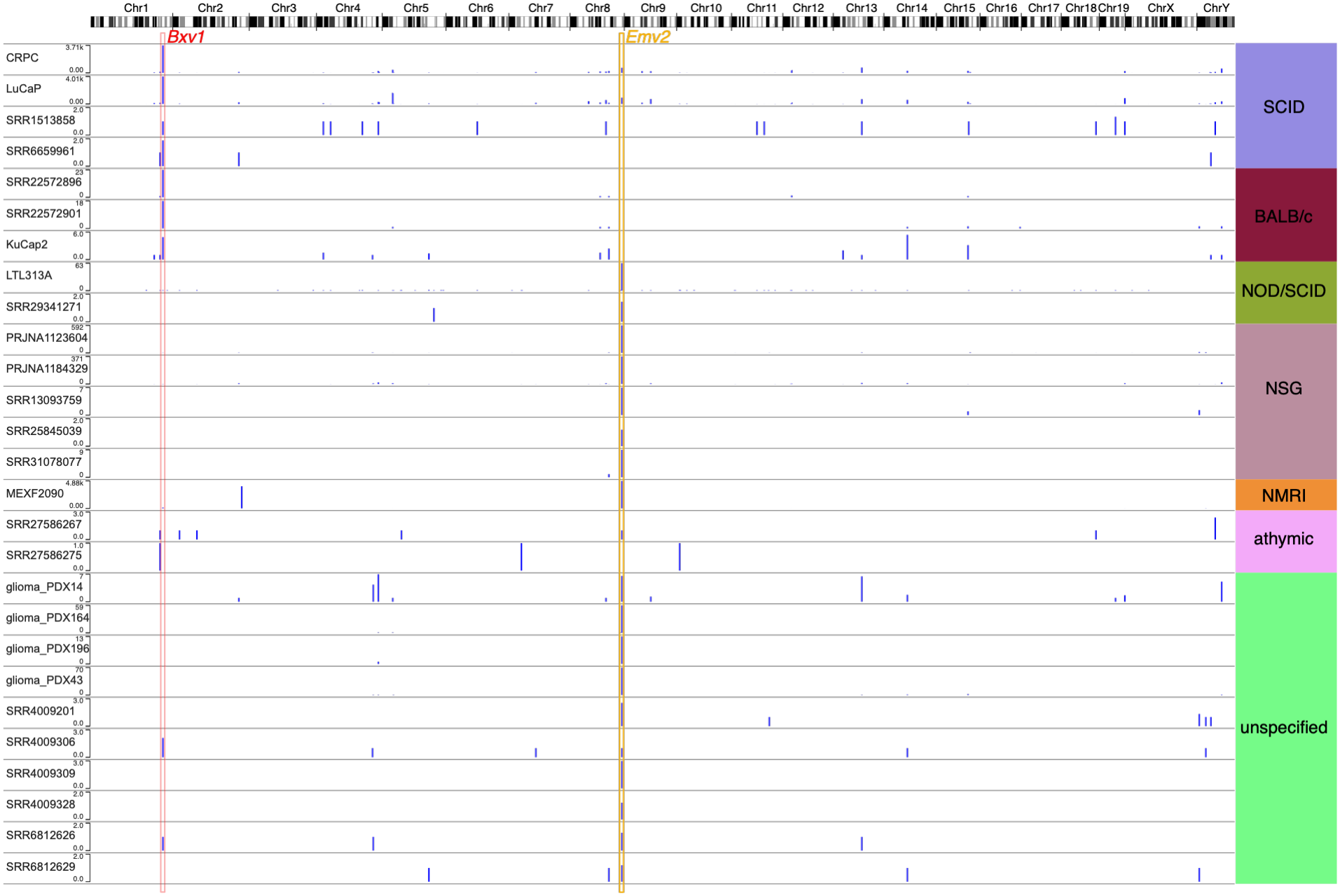
The WashU Epigenome Browser view of human-MLV chimeric reads piled up to the mouse reference genome mm10. All chromosomes were shown horizontally. Each track displays one of the 27 samples, grouped by the mouse strains shown on the right. The high peaks piled up to the provirus *Bxv1* and *Emv2* on mm10 were highlighted with colored boxes.

However, chimeric reads derived from PDX models engrafted in NOD/SCID, NSG, or NMRI strains were found to pile up on *Emv2* on Chromosome 8, the sole E-MLV provirus in C57BL/6J (39). NOD/SCID and NSG strains lack the highly active *Bxv1* provirus and are generally considered safe from X-MLV infection (8). Our observation suggests that PDX models derived from NOD/SCID or NSG, which are *Bxv1*-negative strains, can still be infected by MLV, potentially E-MLV.

### The integration of E-MLV in human genome from melanoma PDX MEXF2090

E-MLV can only infect rodent cells (4, 11). To our knowledge, there has never been a report of E-MLV infecting human cells. To gain insights into the apparent integration of E-MLV into the human genome, we obtained a snap-frozen sample of the melanoma PDX model MEXF2090 directly from vendor, which exhibited a significant number of chimeric reads piled up on *Emv2* (Fig. 3) (11). To characterize the complete MLV insertion, we sequenced its genome using HiFi long reads (Materials and Methods).

Consistent with the presence of mouse cells in the model, we discovered that ∼18% of all MEXF2090 HiFi reads can be classified as mouse-only (Materials and Methods).

After excluding them, we aligned the remaining reads to the human reference genome. By examining the human-MLV chimeric reads from the alignment (Materials and Methods), we identified one MLV integration event with a 4bp TSD at Chromosome 3 in the CHM13 reference genome supported by four HiFi reads (Fig. 4A). We successfully validated the insertion by PCR and sanger sequencing (Fig. 4B). HiFi sequencing also revealed that this provirus was hypomethylated throughout (Fig. 4C). Finally, we generated the complete MEXF2090 MLV provirus from the consensus of these reads and aligned them to X-MLV, P-MLV, and E-MLV using Needleman–Wunsch global pairwise alignment (40). The sliding-window identity between them across the genome shows that the MEXF2090 MLV is distantly related to X-MLV and P-MLV but shares 99.9% identity with E-MLV across its entire genome (Materials and Methods) (Fig. 4D). These data support positive identification of intact E-MLV integration into the human genome in the melanoma PDX MEXF2090.

**Fig. 4:**
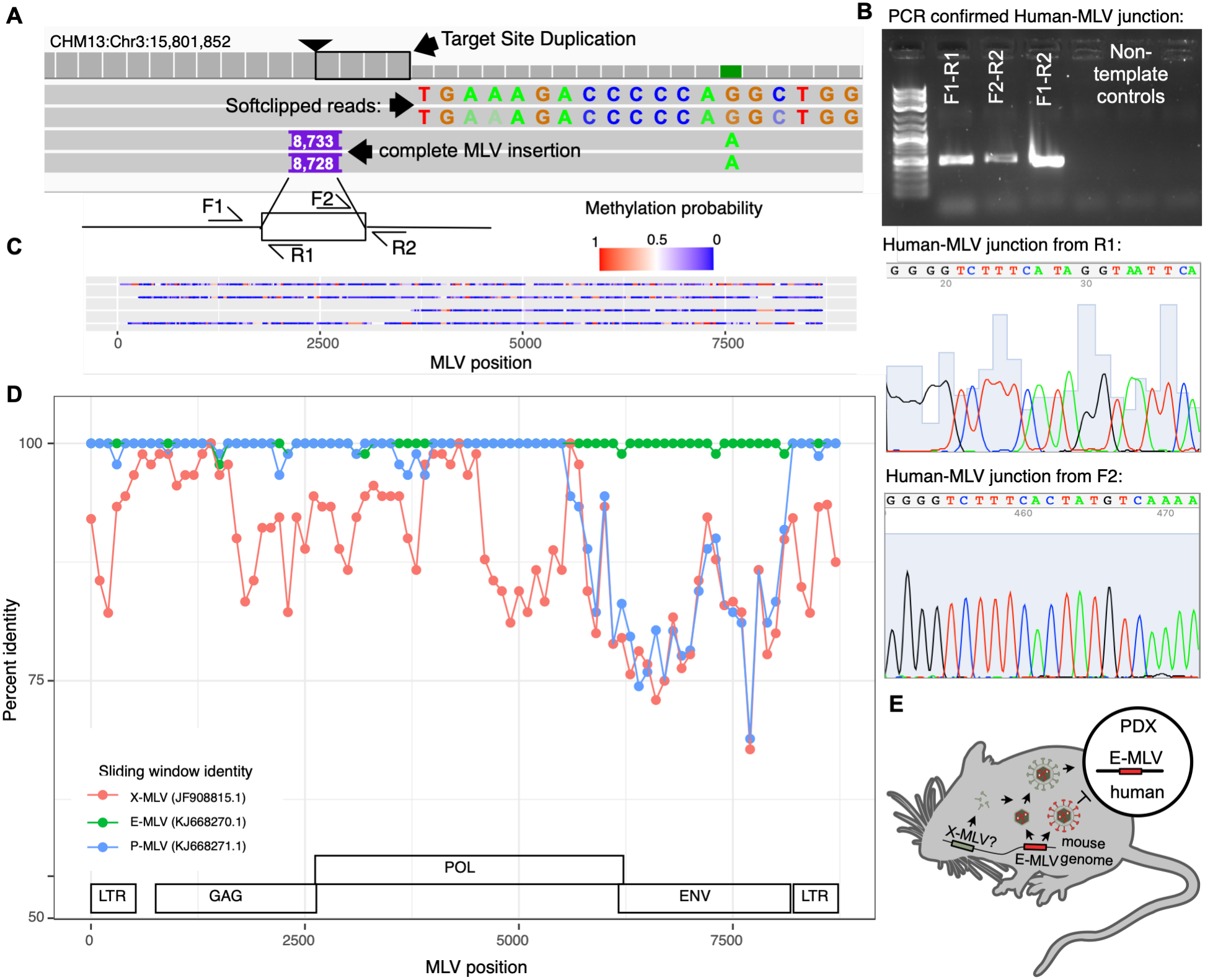
HiFi long-reads support the E-MLV integration in the human genome. (**A**) The IGV screenshot showing the four reads supporting the intact E-MLV integration in the human genome. There are two HiFi reads containing part of the MLV as soft-clip, and two reads containing the whole MLV integration with 4bp TSD. The insertion site TSD, soft-clipped reads match the end of MLV, and the insertions in both reads representing MLV and TSD were annotated. The four PCR primers used for verification were illustrated at the bottom. (**B**) PCR and sanger sequencing validation of the presence of MLV insertion. On the top agarose gel image, each lane from left to right represents PCR product generated from: DNA size ladder, left junction of MLV insertion, right junction of MLV insertion, allele without MLV insertion, and three non-template controls without DNA sample with primer pairs in the same order as lane 2-4. The bottom two panels show the sanger sequencing peaks at human-MLV junctions initiated from primer R1 and F2, respectively. (**C**) Methylation probability of each CpG site on the four supporting reads aligned to the MEXF2090 MLV sequence. The methylation probability was shown as a blue-red color scale. (**D**) Red, green, and blue lines displaying sliding window identity between MEXF2090 and X-MLV, E-MLV, and P-MLV, respectively. MLV structure features were plotted at the bottom. (E) Schematic display of the possible scenario of how E-MLV crossed the species boundary. The E-MLV particle with its own envelope protein can’t infect human cells, but the incorporation of another compatible envelope protein (possibly expressed from a defected X-MLV) makes it possible to infect xenografted human cells.

## Discussion

MLV, a well-studied mouse endogenous retrovirus, has been historically classified into ecotropic, xenotropic, and polytropic varieties based on their host infectivity range (4, 11). Among these, X-MLVs have garnered the most interest due to their potential to infect human cells, culminating in the infamous false alarm of XMRV as a human pathogen (12, 14).

While XMRV and *Bxv1* have been reported to infect human cells through xenograft or indirect contamination, it is generally believed that such cases are rare events and almost certainly do not mediate pathology. X-MLV integration in PDX models is similarly thought to be rare (3, 7). However, in this study, we applied Kmeric to systematically screen all publicly available paired-end PDX genomic reads to discover human-mouse chimeras and found that approximately 25% of the sequencing runs we examined showed a significant enrichment of MLV among human-mouse chimeric reads. Additionally, we discovered that about 90% of all reads spanning the human-MLV junctions in the human genome matched the MLV boundary motif, and 14 out of the 27 human-MLV chimera-enriched samples had canonical MLV TSDs. The identification of indisputable hallmarks of MLV integration in the human genome strongly supports our observations of *bona fide* MLV integrations in human cells.

We discovered that a significant portion of PDX models were infected by MLVs. However, our study does not invalidate all studies based on these PDX models, as we have not demonstrated that these integrations cause functional or phenotypic changes in cells containing them. Nevertheless, we believe it is crucial to characterize these MLV infection events to gain a deeper understanding of their impact on xenografted tumors. To stringently mitigate the possibility of MLV integration confounders on future xenograft studies, we recommend the generation of novel immunocompromised mouse strains with CRISPR knockout of endogenous MLV sequences.

In addition to prevalent MLV integration, we also unexpectedly revealed integration of an E-MLV in the human genome, which was subsequently validated by PCR and Sanger sequencing. The extracellular loop 3 domain (ECL3) of the murine cationic amino acid transporter 1 (CAT1) mediates the selective entry of E-MLV into murine cells, whereas sequence divergence present in human CAT1 ECL3 completely abolishes E-MLV envelope binding (42). To account for the unexpected detection of E- MLV integration, we examined both the receptor-binding domain (RBD) of the MLV envelope and the coding sequence of CAT1 in MEXF2090 but found no amino acid substitutions that would support a novel virus–receptor interaction. We speculate that E-MLV infected MEXF2090 cells by adopting another compatible envelope protein (likely from a defective X-MLV), a process known as pseudotyping (Fig. 4E) (43, 44). If that is the case, we speculate that the integrated MLV provirus in the human genome cannot propagate further infections in human cells alone unless complemented by other envelope proteins expressed from the same cell. Alternatively, it may lose its ability to infect other cells and transform into a retrotransposon (45, 46). E-MLV can also recombine with other retroviruses to create human-compatible infectious particles as previously demonstrated (15, 47).

We found that the risk of MLV infection is not confined to X-MLV: E-MLV can also act as a cross-species transmission agent. It has been speculated that without the most active X-MLV, *Bxv1*-negative strains used in xenograft should be safe from MLV infections (8); however, our results challenges this assumption. Numerous PDX samples derived from *Bxv1*-negative strains were infected by MLV, with E-MLV as the most likely culprit (Fig. 3). HiFi and Sanger sequencing further support integration of E-MLV in the genome of melanoma PDX MEXF2090 (Fig. 4).

These data underscore the heightened risk of cross-species viral transmission in xenografts. Here we investigated viral transmission between mouse and human using abundantly available PDX sequencing data. In addition to PDX oncology models, many other murine xenograft models are widely used in medical research such as CD34+ stem cell humanization for immunology and ectopic organs for regenerative medicine (2, 48), and they may be similarly susceptible to MLV infection. Accordingly, scientists should attempt to characterize the potential for infection, integration and functional interference in their models.

Beyond mouse-to-human transmission, it is not difficult to envision similar cross-species infections between other species. One particularly concerning area impacted by our findings is the pig-originated porcine endogenous retrovirus (PERV) in the context of human xenotransplantation. Similar to MLV, PERV is an endogenous retrovirus capable of infecting human cells *in vitro*. PERV transmission during xenotransplantation poses a potential risk not only to transplant recipients but also to the general public should the virus acquire the capacity for sustained human-to-human transmission. Notably, CRISPR-based strategies have been developed to inactivate all genomic copies of PERV in the pig genome (9, 10, 49), and several recent xenotransplantation studies have used these PERV-inactivated donor strains, although others have continued to use conventional pigs (50–54). Given the prevalence and unexpected nature of the MLV infections identified in our study, we recommend rigorous validation to ensure that porcine donor tissues are truly free of transmissible PERVs. Furthermore, our results demonstrate our pipeline Kimeric as an efficient approach for detecting viral integration in the human genome, which could serve as a valuable tool for monitoring xenotransplantation safety.

Finally, our study highlights the importance of funding and maintaining public accessible research data and open-source software. In addition to enabling community-based validation to improve reproducibility, these data could also inspire new research and applications that were not anticipated initially, while tools developed for metagenomics and 3D genome can be adopted to detect the integration of viruses in the human genome. We hope our study can inspire more novel applications of existing genomic datasets and methodologies.

## Materials and Methods

### Kraken2

We downloaded Kraken2 from GitHub (https://github.com/DerrickWood/kraken2) and build a custom Kraken2 database using only human and mouse reference genomes (31). With Kraken2 installed and the custom database built, we ran Kraken2 classifier on each end of the fastq files separately and extracted only pairs with one end classified as human and the other end classified as mouse. At last, we removed pairs if at least 50% of one of the ends was masked as low complexity by pydustmasker (32).

The pipeline (kraken2_install.sh and kraken2_run.sh) are provided in the GitHub repo (https://github.com/xzhuo/kmeric).

### Alignment of human-mouse chimeric reads

We filtered out low-quality reads using fastp from Kraken classified human-mouse chimeric reads. We then aligned high quality paired-end human-mouse chimeric reads to the combined human-mouse reference genome using ‘bwa men -5SP’ option to ensure the separated mapping of each end (34). After alignment, we removed mapping duplicates using Picard (‘picard MarkDuplicates’) (https://broadinstitute.github.io/picard/). We converted the alignment bam file to paired- end bedpe file using BEDTools (‘bedtools bamtobed -bedp’) and filtered out pairs with both ends mapped to the same species (36). From the ‘bedpè file we generated one ‘bed’ file including human coordinates and one ‘bed’ file with mouse coordinates. At last, we ran the Fisher’s exact test using the mouse bed file of chimeric reads and RLTR4_Mm coordinates generated from UCSC repeatMasker output. Based on the test, we selected 397 sequencing runs with enrichment ratio > 9 and P-value < 1e-09 as extremely enriched with human-MLV chimeric reads. The alignment pipeline is provided as a Snakefile from the GitHub repo (https://github.com/xzhuo/kmeric).

### Softclip motif

From the deduplicated bam file we extracted human-MLV chimeric reads from samples where they are highly enriched. We then separated the human ends and the mouse ends to two different bam files. From the bam file including all the ends mapped to human, we ran a Python script ‘scripts/extract_bam_softclip.py’ from the GitHub repo (https://github.com/xzhuo/kmeric) to define each read by whether it begins the soft clipped end with motif ‘TGAAÀ. We ran another Python script ‘ extract_bam_softclip_woadj.py’ from the same repository to extract the aligned sequences in the mouse genome (the supplementary alignment if the primary alignment is in human and the primary alignment if the supplementary alignment found in human), and aligned them to the RLTR4_Mm consensus using bwa-mem (34).

### Map human-MLV chimeric reads to the mouse genome

From the bam file containing all human-MLV chimeric reads mapped to the mouse reference genome mm10, we created bigwig files from reads pileup through ‘genomecov’ in BEDTools and ‘bedGraphToBigWig’ in UCSC Kentutils and made them available at (https://epigenomegateway.wustl.edu/browser2022/?sessionFile=https://wangcluster.wustl.edu/∼xzhuo/mlv/bigwig/all27.mm10.session.json).

### TSD identification

We grouped all samples with human-MLV chimeric enriched to 27 samples, and aggregated all reads mapped to the human genome from the same sample to a single bed file. We then manually examined all loci with reads mapped to both strands for potential target site duplications.

### HiFi sequencing

We obtained snap-frozen MEXF2090 sample from Charles River and extracted high molecular weight DNA using the standard protocol and sequenced the DNA library using PacBio Revio platform. The HiFi reads were uploaded the NCBI SRA (PRJNA1466899). We characterized CpG methylation of HiFi reads using Jasmine V26.1.2 (https://github.com/PacificBiosciences/jasmine).

To classify HiFi reads, we converted HiFi reads from bam format to fastq format and ran Kraken2 with the same custom human-mouse database (31). We excluded all HiFi reads classified as mouse and contains <1000 cumulative human Kmers. We then aligned the rest of all reads to the human reference genome CHM13 using minimap2 (55). We also Blast searched human-mouse chimeric reads against MLV sequence to identify potential human-MLV chimeric (56). At last, from the alignment of human-MLV chimeric reads to the human reference genome we identified four reads (m84081_260329_053816_s1/225773966/ccs, m84081_260329_053816_s1/86902467/ccs, m84081_260329_053816_s1/200214583/ccs, m84081_260329_053816_s1/87165854/ccs) supporting the MLV integration event in the human genome.

### Methylation of MEXF2090-MLV

We extracted the soft clipped and inserted sequences from the four reads using script ‘scripts/bam.extractInsertion.py’. We then attached the methylation tags to the four truncated reads using ‘modkit repair’. We aligned the four truncated reads using MUSCLE and generated the MEXF2090-MLV provirus sequence from the consensus of the multiple sequence alignment (57). After aligning each of the truncated reads to the MEXF2090-MLV using minimap2, we extracted the methylation of each CpG site using ‘modkit extract full --cpg’ from the alignment (https://nanoporetech.github.io/modkit/).

### Alignment of MEXF2090-MLV to X-MLV, E-MLV, and P-MLV

We also aligned the MEXF2090-MLV sequence to each of the X-MLV (JF908815.1), E- MLV (KJ668270.1), and P-MLV (KJ668271.1) using the Needleman-Wunsch pairwise Global Alignment and calculated the 100bp sliding window identity using Perl script ‘scripts/sliding_window.aln.pl’.

### PCR validation and sanger sequencing of MEXF2090-MLV

Based on the MEXF2090-MLV integration site on CHM13 Chromosome3 we designed PCR primers to amplify both junctions of the inserted MLV. The four primers listed in Fig. 4 are F1: 5’-CGGCTTAACTGCTTTCGTGT-3’, R1: 5’- GATCCTTGGGAGGGTCTCCT-3’, F2: 5’-GAGGAGACCCTCCCAAGGAT-3’, and R2: 5’-ATTACCCCGTGCCCTTTTGA-3’, Primer R1 and F2 were used to validate the PCR products by sanger sequencing.

## Data and code availability

All data are available in the main text or the supplementary materials, and the kmeric pipeline is available at https://github.com/xzhuo/kmeric.

## Acknowledgments

Here we want to thank Georg Kuales, Torsten Giesemann, and Tobias Deigner from Charles River Germany for facilitating the shipment of the MEXF2020 sample.

Our work is supported by: National Human Genome Research Institute grant R01HG007175 (TW) National Human Genome Research Institute grant U41HG010972 (TW) National Human Genome Research Institute grant U41HG010971 (TW) National Human Genome Research Institute grant U24HG012070 (TW) National Institute of Neurological Disorders and Stroke grant U24NS132103 (TW) National Institute on Drug Abuse grant UM1DA058219 (TW)

